# SOPHIE: viral outbreak investigation and transmission history reconstruction in a joint phylogenetic and network theory framework

**DOI:** 10.1101/2022.05.05.490757

**Authors:** Pavel Skums, Fatemeh Mohebbi, Vyacheslav Tsyvina, Pelin Icer Baykal, Alina Nemira, Sumathi Ramachandran, Yury Khudyakov

## Abstract

Genomic epidemiology is now widely used for viral outbreak investigations. Still, this methodology faces many challenges. First, few methods account for intra-host viral diversity. Second, maximum parsimony principle continues to be employed, even though maximum likelihood or Bayesian models are usually more consistent. Third, many methods utilize case-specific data, such as sampling times or infection exposure intervals. This impedes study of persistent infections in vulnerable groups, where such information has a limited use. Finally, most methods implicitly assume that transmission events are independent, while common source outbreaks violate this assumption.

We propose a maximum likelihood framework SOPHIE (SOcial and PHilogenetic Investigation of Epidemics) based on integration of phylogenetic and random graph models. It infers transmission networks from viral phylogenies and expected properties of inter-host social networks modelled as random graphs with given expected degree distributions. SOPHIE is scalable, accounts for intra-host diversity and accurately infers transmissions without case-specific epidemiological data. SOPHIE code is freely available at https://github.com/compbel/SOPHIE/

## 1 Introduction

Continuing advances of sequencing technologies had a profound effect on virological and epidemiological research. In particular, they vitalized *genomic epidemiology* – an interdisciplinary area of research that uses analysis of viral genomes to understand how viruses evolve and spread. As a result, methods of genomic epidemiology are becoming major tools for investigation of outbreaks and surveillance of transmission dynamics [2, 5]. It became possible largely due to the rapid progress in development of efficient computational methods. The list of transmission history inference tools published over the last decade includes Outbreaker and Outbreaker 2 [39, 10], SeqTrack [38], SCOTTI [19], Phybreak [41], Bitrugs [75], BadTrIP [18], Phyloscanner [76], StrainHub [17], TransPhylo [22], STraTUS [35], TreeFix-TP [68], QUENTIN [67], VOICE, HIVTrace [43], GHOST [46], MicrobeTrace [7], SharpTNI [64], TiTUS [65], TNeT [21] and others [78, 48, 23, 49, 16, 9, 34]. These tools have been successfully applied to HIV, hepatitis C virus (HCV), severe acute respiratory syndrome (SARS), Middle East respiratory syndrome (MERS), severe acute respiratory syndrome coronavirus 2 (SARS-CoV-2) and other viruses [74, 59, 79, 57, 42, 8].

The hallmark of viruses as species is an extremely high genomic diversity originating from their error-prone replication. As a result, each infected individual usually hosts a heterogeneous population of numerous genomic variants. Such populations are traditionally called *viral quasispecies* [24]. First generation of transmission inference methods largely ignored intra-host viral diversity and considered only a single sequence per host (usually consensus). Later, it has been demonstrated that taking intra-host diversity into account greatly enhances the predictive power of transmission inference algorithms [76, 67, 1, 61, 42]. In particular, it allows to detect the viral evolution directionality in situations when a reliable phylogenetic rooting is not possible [67, 61, 30] – such situation is very common for HIV, HCV and other long-standing epidemics, as well as for the regional epidemics of SARS-CoV-2 characterized by multiple introductions of the virus. First phylogenetic approaches to infer transmission directions using viral quasispecies appeared independently and almost simultaneously in [61] and [30]. Later, the ideas of [61] have been incorporated into the full transmission network inference tool Phyloscanner [76], while the methodology of [30] has been utilized by QUENTIN [67]. These tools were followed by TNeT [21], TiTUS [65], SharpTNI [64], BadTrIP [18], all of which are specifically tailored to take into account intra-host viral diversity.

Despite the significant progress achieved with the appearance of the next generation of transmission inference method, a number of computational, modelling and algorithmic challenges still need to be addressed.

1. Most new tools utilize a maximum parsimony principle and many of them are based on various extensions of the classical Sankoff labelling algorithm. However, maximum likelihood or Bayesian phylogenetic models are richer and incorporate additional inferred temporal information that can be used for more accurate reconstruction of transmission links [51].
2. Several studies demonstrated that in many cases genomic data alone do not allow to resolve ambiguities in transmission network inference, and so the incorporation of additional evidence is necessary [37, 72, 39]. Such evidence most often comes in the form of case-specific epidemiological information. However, most common types of such information are useful only in particular settings. For example, many tools use sample collection times to identify the order of infections. However, HIV, HCV and many other infections tend to be initially asymptomatic, and consequently, sampling times may not accurately reflect the actual infection times. Other tools rely on exposure intervals for the same purpose. However, in outbreaks with high transmission rates (e.g. in HIV/HCV outbreaks associated with injection drug use or during the global pandemic of SARS-CoV-2/Influenza), many susceptible hosts are almost constantly exposed to the virus, thus effectively making exposure intervals useless.
3. Most methods implicitly assume that transmission network edges are independent. Such assumption is associated with *random mixing* models, that suppose that differences between individuals are negligible and any person can infect any other person with the same probability. However, this is not always the case, as, for example, certain hosts infect more people than an average individual [29].
4. The models that use more comprehensive and parameter-rich models lead to computationally hard optimization problems. To find transmission networks and estimate other parameters, such methods mostly rely on Markov Chain Monte Carlo (MCMC) sampling from the model parameter space [18, 67, 19]. Given that the parameter spaces are enormous [35], such strategy is computationally expensive and may produce sub-optimal results.

In this study, we propose to address these challenges by integrating phylogenetic and random graph models. Our major idea is to bring into consideration the social component of the epidemics. Infectious diseases spread over the social networks of contacts between susceptible individuals, and transmission networks to a significant degree mirror the properties of these social networks [44, 73, 36, 60]. These properties are well defined in network theory, sociology and classical epidemiology [52]. In light of this, we propose to infer transmission networks by integrating two components: the evolutionary relationships between viral genomes represented by their phylogenies and the expected structural properties of inter-host social networks. Frequently cited properties of social contact networks include power law degree distribution, small diameter, modularity and presence of hubs [3, 52]. All of them are reflected by network vertex degrees. Thus, we model social networks as random graphs with given expected degree distributions (EDDs). They are commonly scale-free [73, 6], but our method can handle more specific EDDs of needle-sharing networks, sexual-contact networks or networks obtained by epidemiological contact tracing or respondent-driven sampling. The goal is to find transmission networks that are consistent with observed genomic data and have the highest probability to be subnetworks of random contact networks.

This methodology is implemented within a maximum likelihood algorithmic framework SOPHIE (SOcial and PHilogenetic Investigation of Epidemics). SOPHIE samples from the joint distribution of phylogeny ancestral traits defining transmission networks, estimates the probabilities that sampled networks are subgraphs of a random contact network and summarize them accordingly into the consensus network. This approach is scalable, accounts for intra-host diversity and accurately infers transmissions without case-specific epidemiological data.

We applied SOPHIE to synthetic data simulated under different epidemiological and evolutionary scenarios, as well as to experimental data from epidemiologically curated HCV outbreaks. The experiments confirm the effectiveness of the new methodology.

## 2 Results

### 2.1 SOPHIE algorithm for inference of transmission networks

We developed SOPHIE - a modelling and algorithmic framework to infer viral transmission networks from genomic data by integrating phylogenetic and random graph models. Within this framework, we define the *transmission network inference* problem as follows. We are given a time-labelled phylogeny *T* = (*V* (*T*), *E*(*T*)) with *n*_*l*_ leafs corresponding to viral haplotypes sampled from *n*_*h*_ infected hosts; each leaf *u* is assigned the label *λ*_*u*_ ∈ [*n*_*h*_] corresponding to its host. Such tree can be constructed using standard phylogenetic tools such as RAxML [70], PhyML [32] and IQ-Tree [53]. The goal is to extend ***λ*** to internal nodes in an optimal way. In this model, every multi-labelled tree edge *uv* corresponds to a direct or indirect transmission between the hosts *λ*_*u*_ and *λ*_*v*_. Thus, the transmission network *G* = *G*(*T*, ***λ***) with the vertex set *V* (*G*) = [*n*_*h*_] can be constructed by contracting the vertices with the same label [34] (Fig. 2). The simplest variant of this problem is the *maximum parsimony label inference* where the goal is to minimize the number of transmission events. It can be easily solved using e.g. Fitch or Sankoff algorithms [63, 28] and their modifications. However, straightforward maximum parsimony approach alone often leads to epidemiologically unrealistic results [76]; furthermore, there are usually many most parsimonious solutions [20, 65]. Within maximum likelihood framework, ancestral labels can be inferred using so-called “migration model” [62]. In this case, Fitch or Sankoff algorithms can be replaced by the dynamic programming algorithm of Pupko et.al. [56] or its extensions [62]. However, as mentioned above, phylogenetic signal alone can be insufficient for accurate transmission network reconstruction [37, 72, 39]. In particular, in the absence of reliable estimations of transmission rates between individual hosts, migration-based approaches have to rely on simple substitution models; as a result, similarly to the case of maximum parsimony, the numbers of near-optimal solutions can be high.

**Figure 1:**
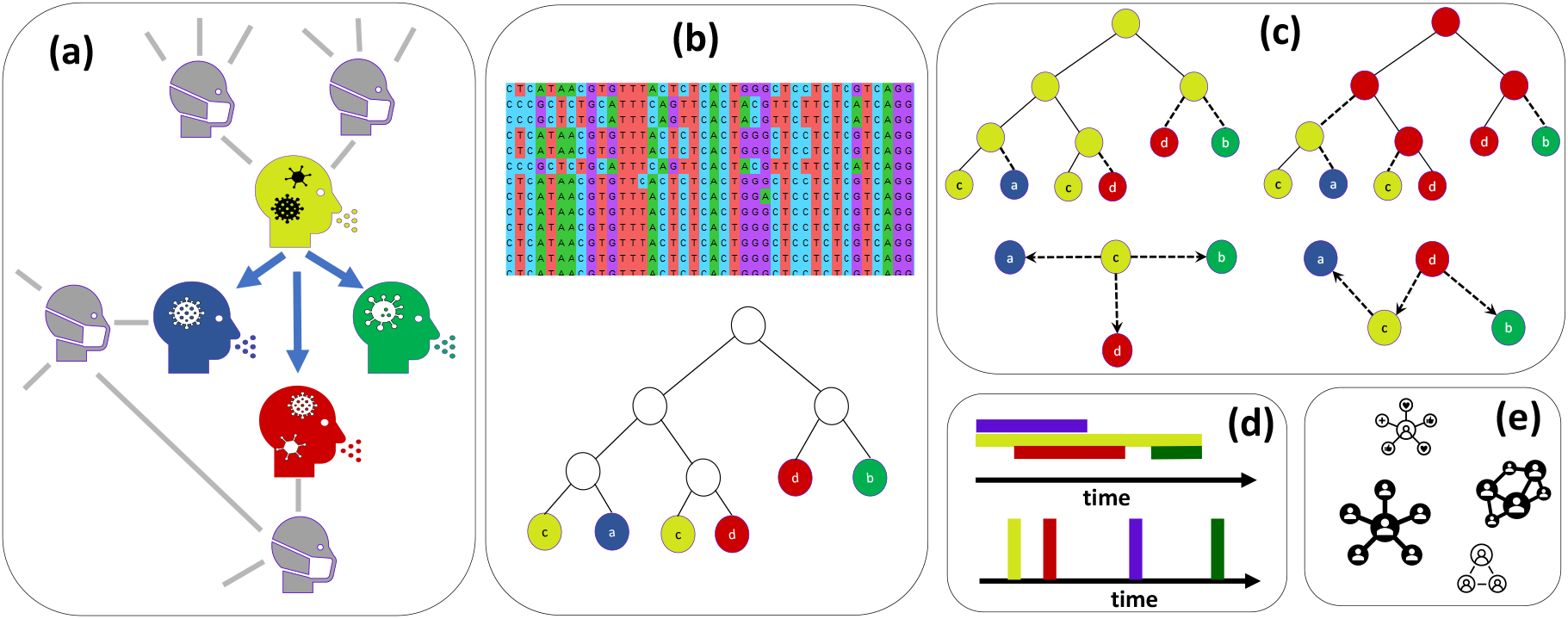
Approaches and challenges for transmission history reconstruction using genomic data. **(a)** *Example of a viral outbreak and its transmission network* consisting of 4 individuals (highlighted in light green, blue, dark green and red) and 3 transmissions links (blue arrows). The transmission network is a part of a larger unobserved social network of contacts between susceptible individuals (the unobserved part is highlighted in gray). Social networks serve as conduits for the infection spread, and thus transmission networks reflect the properties of social networks. Due to the high virus mutation rates, each infected individual hosts a population of related but distinct viral genomic variants (viral quasispecies). **(b)** *First step of genomic epidemiology investigation*. Intra-host viral variants are sequenced, de-noised and aligned; the obtained viral haplotypes are used to construct a viral phylogeny. Leafs of this phylogeny correspond to sampled viral variants and labelled by their hosts (colors of the leafs correspond to the colors in (a)). **(c)** *Phylogenetic inference of transmission networks*. Labels of leafs are extended to internal nodes, and every tree edge with multi-labelled end nodes defines a transmission between the corresponding hosts. Two possible ancestral label assignments are depicted. Tree edges defining transmissions are dashed, the corresponding transmission network is shown below each assignment. Note that both assignments have the same number of such edges, i.e. the same parsimony score. Thus, parsimony does not allow to rank the obtained transmission networks. **(d)** *Resolution of phylogenetic ambiguities using case-specific epidemiological data* proposed in prior studies. One possibility is to consider patient exposure intervals (upper figure): in this example the intervals for the red and green patients do not overlap, thus ruling out the second network containing a link between these patients. Another possibility is to take into account sampling times (lower figure): light green patient was sampled earlier thus making more probable the first network, where it is a root. Unfortunately, such information often has a limited use for many real outbreaks of HIV, HCV, SARS-CoV-2, etc. **(e)** *Resolution of phylogenetic ambiguities using the prior knowledge about social network properties*. We propose to integrate phylogenetic and random graph models: first, we sample transmission networks from the phylogeny-based distribution, and then measure their agreement with expected properties of the distribution of inter-host social networks. In this example, the depicted social network distribution favors the first candidate transmission network that has more “star-like” structure.

**Figure 2:**
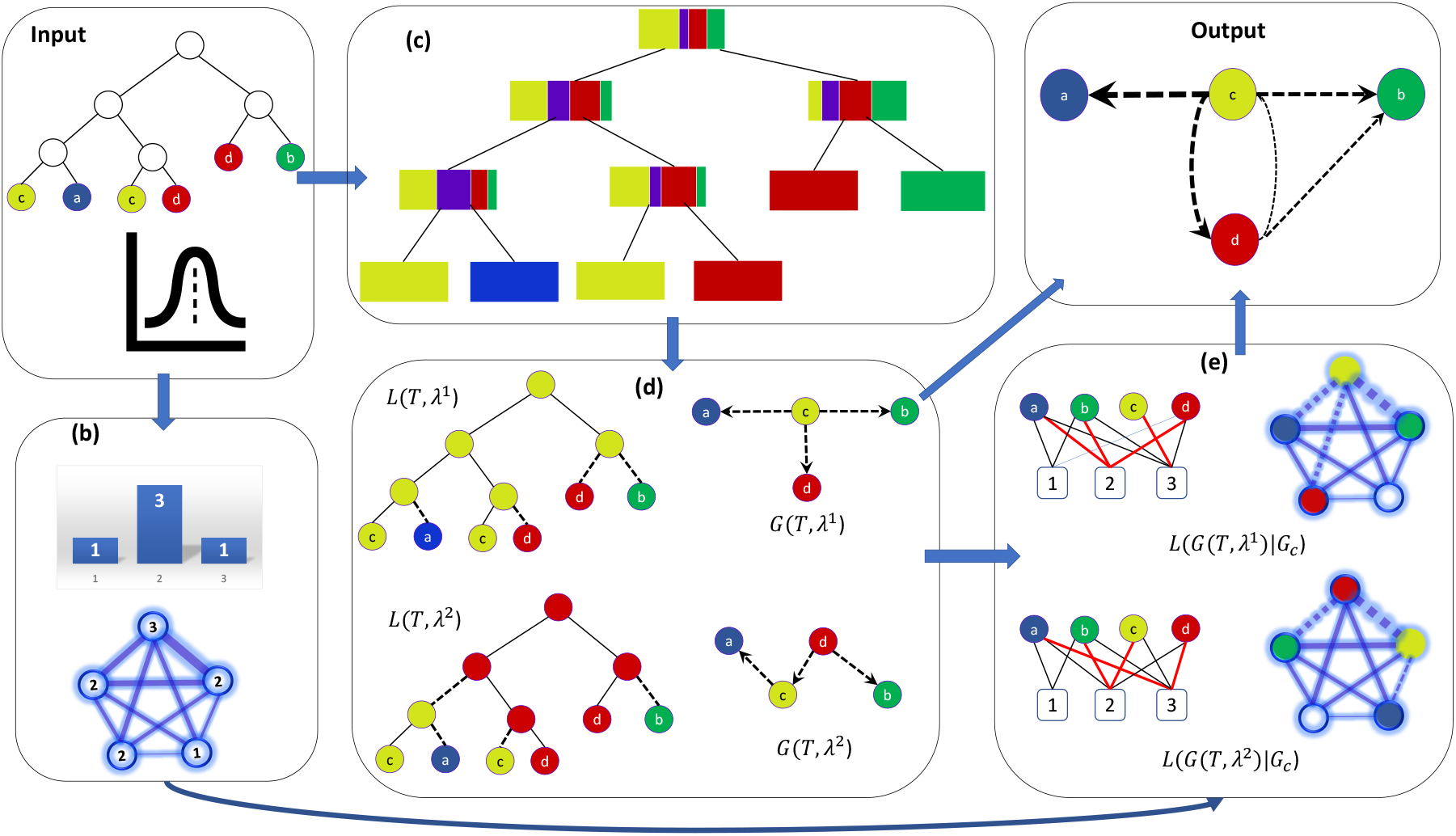
Joint phylogenetic and random graph-based approach for transmission history reconstruction implemented in SOPHIE. Input: a labelled phylogeny with leafs corresponding to viral haplotypes from 4 infected hosts (highlighted in different colors); expected degree distribution of a contact network that contains the true transmission network as a subgraph. **(b)** Generalized Random Graph (GRG) model of a contact network depicted as a complete graph with edge thicknesses proportional to their probabilities. It is accompanied by the expected degree counts of contact network vertices. **(c)** SOPHIE samples from the joint distribution of ancestral labels assignments using a dynamic programming. First, the algorithm performs a post-order traversal and calculates, for each internal node, conditional likelihoods of observing the labels of its descendants given a label of this node. On a figure, the widths of colored strips are proportional to the conditional likelihoods given the hosts with the corresponding color-codes. After all conditional likelihoods are calculated, the algorithm performs a pre-order traversal and samples a label for each node from the corresponding posterior distribution given its parent’s sampled state (see Subsection 4.1.1). **(d)** Two sampled ancestral label assignments *λ*_1_ and *λ*_2_, the corresponding transmission networks and their phylogenetic likelihoods. Tree edges defining transmissions are dashed. The networks are obtained by contracting the tree nodes with the same labels. **(e)** SOPHIE calculates network likelihoods of sampled transmission networks by embedding them into random contact networks. To find an embedding, SOPHIE maps the transmission network vertices to their degrees in the contact network. It is done via the reduction to a generalized uncapacitated facility location problem with convex costs, where the hosts serve as clients and their possible expected degrees in – as facilities. On the left side of the panel, the instances of the facility location problem for two sampled networks are depicted. Optimal client assignments are highlighted in red, next to them the corresponding embeddings of transmission networks into contact networks are shown. See Subsection 4.1.2 for details. **Output:** a consensus of sampled transmission networks. Edges represent possible transmission links, their thicknesses are proportional to their inferred likelihood supports. See Subsection 4.1.3 for details.

In light of this, we extend a maximum likelihood approach by integrating a phylogenetic model with a model of social networks of susceptible individuals. Under this methodology, a transmission network is defined by two properties: it is a contraction of the phylogeny and, at the same time, a subgraph of a inter-host contact network of susceptible individuals (Fig. 2). In reality, the contact network is not directly observed. Therefore, we model it as a random graph with the *expected degree distribution (EDD)*. EDD carries information about structural and spectral properties of contact networks [12, 52, 13], and can be adjusted to reflect specific epidemiological settings.

The general scheme of our approach is as follows (Fig. 2):

1. We consider phylogeny node labels as discrete traits and sample from the joint distribution of label assignments under the selected substitution model (Subsection 4.1.1).
2. For each sampled label assignment ***λ***, we construct the corresponding transmission network *G*(*T*, ***λ***) and estimate its *network likelihood*, which is defined as the maximum probability that this network is a subgraph of a random contact network with the given EDD (Subsection 4.1.2)
3. Estimate the final transmission network as a weighted consensus of sampled networks. The edge weights here represent the inferred joint likelihood network-based and phylogeny-based likelihood support for the corresponding transmission links.

Each of these steps is described in detail in Methods section 4.

### 2.2 Algorithm benchmarking

We validated SOPHIE on synthetic and experimental data with known transmission networks. To evaluate the accuracy of inferred networks, we estimated sensitivity (i.e. the fraction of inferred transmission edges among true transmission edges), specificity (i.e. the fraction of true transmission edges among inferred transmission edges) and *f* -score (i.e. the harmonic mean of sensitivity and specificity). The latter parameter has been used as the principal evaluation metric.

In this study, SOPHIE was compared with Phyloscanner and TNet. Both methods are based on maximum parsimony principle: Phyloscanner reconstructs ancestral labels using a Sankoff algorithm with specially adjusted parsimony scores, while TNet uniformly samples from the space of most parsimonious label assignments that minimize the number of back transmissions. Other published phylogeny-based tools that account for intra-host viral diversity, TiTUS and BadTrIP, utilize case-specific exposure intervals as an additional source of information. Theoretically, in the absence of exact exposure dates, both tools can work with arbitrarily large exposure intervals. However, as noted by the authors of BadTrIP [18], such assumption has a significant negative effect on in the accuracy of their method. We observed the similar effect for TiTUS: its average *f* -score was quite low (mostly within a range of ∼ 0.10 − 0.20), thus suggesting that non-trivial exposure intervals are essential for it. Therefore, for the sake of fairness TiTUS and BadTrIP were excluded from further comparison.

#### 2.2.1 Simulated data

To generate synthetic data, we used FAVITES [50] – a flexible tool that can simultaneously simulate viral sequences, phylogenies, contact networks and transmission networks under different evolutionary and epidemiological scenarios. In our case, we assumed that the virus spread over a contact network of 100 susceptible individuals generated using the Barabasi-Albert model [3]. Transmission networks and data sampling were simulated under two epidemiological scenarios:

E1) Susceptible-Infected (SI) transmission model and simultaneous sampling of all infected individuals at the end of the simulation. This scenario corresponds to the typical settings of HIV or HCV outbreaks [57, 54].

E2) Susceptible-Infected-Recovered (SIR) transmission model, with each individual sampling time being chosen from its infection time window. This scenario describes epidemics and surveillance of Influenza, SARS-CoV-2 and other viruses that are associated with acute rather than chronic infections.

Inside each host, viral phylogenies evolved under a coalescent model with two effective population size growth modes:

I1) Exponential effective population growth.

I2) Logistic effective population growth.

For each of four combinations of scenarios E1-E2 and I1-I2, 100 simulated datasets have been generated, with 10 genomes sampled per infected host. For each dataset, we applied SOPHIE, Phyloscanner and TNet to two trees: a true phylogeny provided by FAVITES and a phylogeny reconstructed by RAxML [70]. For a network likelihood calculation with SOPHIE, we used a power-law distribution as an expected degree distribution. In this case, the algorithm has a power-law degree exponent *α* as a hyperparameter. We analyzed SOPHIE performance with the best exponent from the interval (1, 2] and with the exponent randomly drawn from the gamma distribution with the mean 1.6. For each test instance, 100, 000 internal label assignments were sampled, and the final network calculated as a maximum-weight arborescence of the consensus network (see Subsection 4.1.3). Further details can be found in Methods section.

The results of SOPHIE evaluation and comparison with other methods are shown in Tables 1-2 and on Figures 3 - 4. First, the value of the exponent *α* does not significantly affect the results. This demonstrates that accounting for the general shape of the expected degree distribution plays the most important role here, while guessing the best exponent allows for a moderate improvement. Second, for all eight experiments (four combinations of scenarios and 2 types of trees), we found that SOPHIE allows for a statistically significant improvement over TNet and Phyloscanner (*p <* 0.05, Kruskal-Wallis test). The average *f* -score of SOPHIE over all datasets is 0.71 (standard deviation 0.17) and can be as high as 0.92 and 0.90 (for SIR transmission models with the exponential and logistic coalescent and the true phylogeny). The average best absolute *f* -score improvement with respect to existing methods were 0.22 (standard deviation 0.09) over TNet and 0.25 (standard deviation 0.07) over Phyloscanner.

**Table 1:** Mean f-scores of SOPHIE, TNet and Phyloscanner for different simulated and real datasets.

|  | True tree |  |  |  | RAxML tree |  |  |  | Real |
| --- | --- | --- | --- | --- | --- | --- | --- | --- | --- |
|  | SIR exp | SIR log | SI exp | SI log | SIR exp | SIR log | SI exp | SI log |  |
| SOPHIE (best $\alpha$) | 0.92 | 0.90 | 0.82 | 0.41 | 0.68 | 0.67 | 0.78 | 0.48 | 0.70 |
| SOPHIE ($\alpha \sim \Gamma$) | 0.89 | 0.86 | 0.75 | 0.33 | 0.63 | 0.61 | 0.73 | 0.42 | |
| TNet | 0.57 | 0.63 | 0.50 | 0.24 | 0.53 | 0.55 | 0.50 | 0.36 | 0.58 |
| PhyloScanner | 0.75 | 0.71 | 0.48 | 0.03 | 0.49 | 0.45 | 0.50 | 0.29 | 0.37 |

**Table 2:** *p*-values of multiple comparison for Kruskal-Wallis test.

|  | True tree |  |  |  | RAxML tree |  |  |  |
| --- | --- | --- | --- | --- | --- | --- | --- | --- |
|  | SIR exp | SIR log | SI exp | SI log | SIR exp | SIR log | SI exp | SI log |
| SOPHIE vs TNet | 1.6e-12 | 1.4e-7 | 2.1e-19 | 7.1e-7 | 3.0e-3 | 4.7e-2 | 8.4e-16 | 4.7e-3 |
| SOPHIE vs Phyloscanner | 1.2e-4 | 6.3e-5 | 4.7e-21 | 0 | 4.2e-4 | 1.2e-4 | 6.3e-16 | 3.0e-6 |
| TNet vs Phyloscanner | 5.3e-3 | 4.4e-1 | 9.3e-1 | 1.3e-13 | 8.6e-1 | 1.9e-1 | 9.9e-1 | 1.9e-1 |

**Figure 3:**
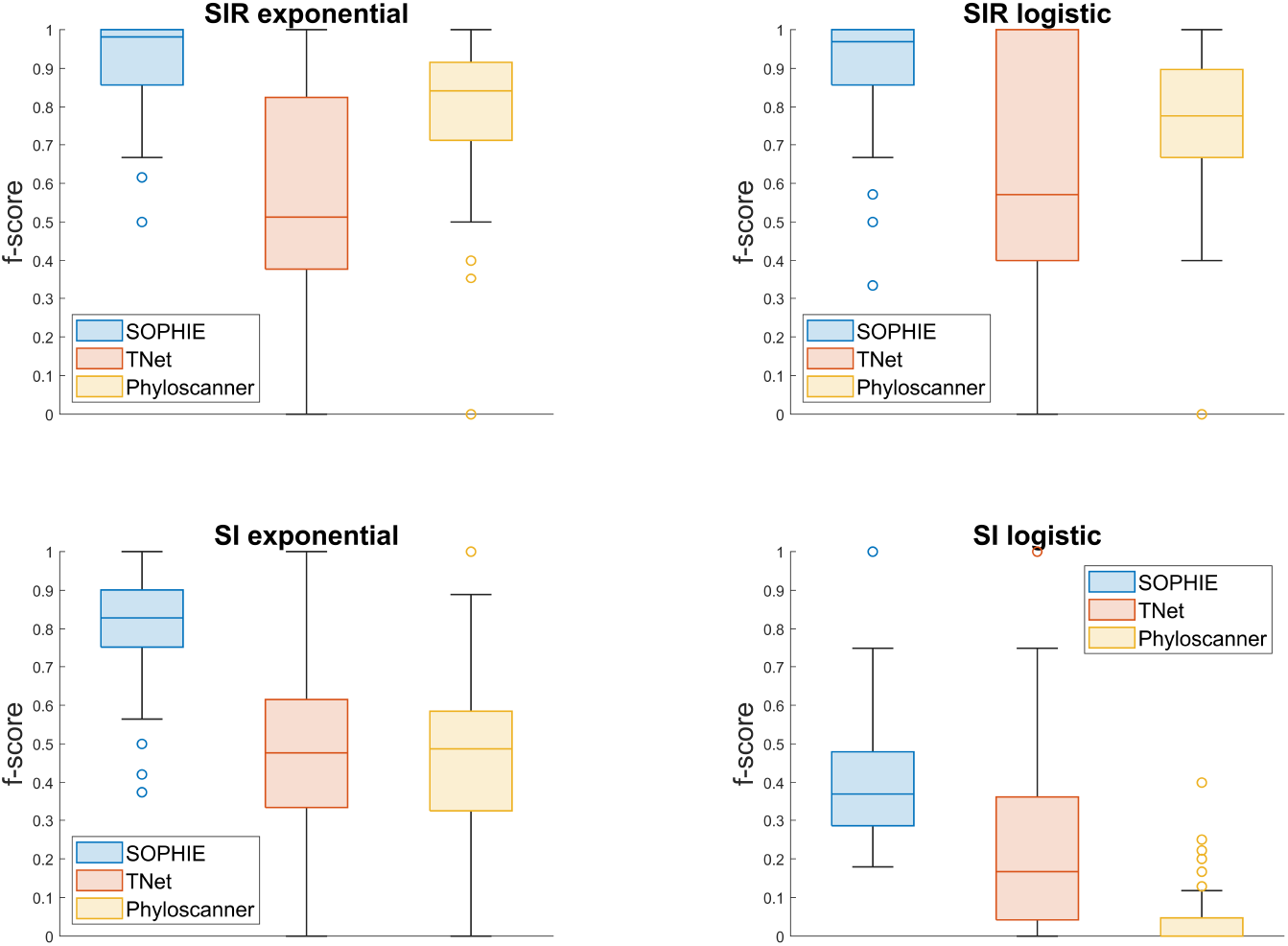
Comparative results of SOPHIE (best exponent), TNet and Phyloscanner on simulated data under different epidemiological and evolutionary scenarios with the true tree simulated by FAVITES

**Figure 4:**
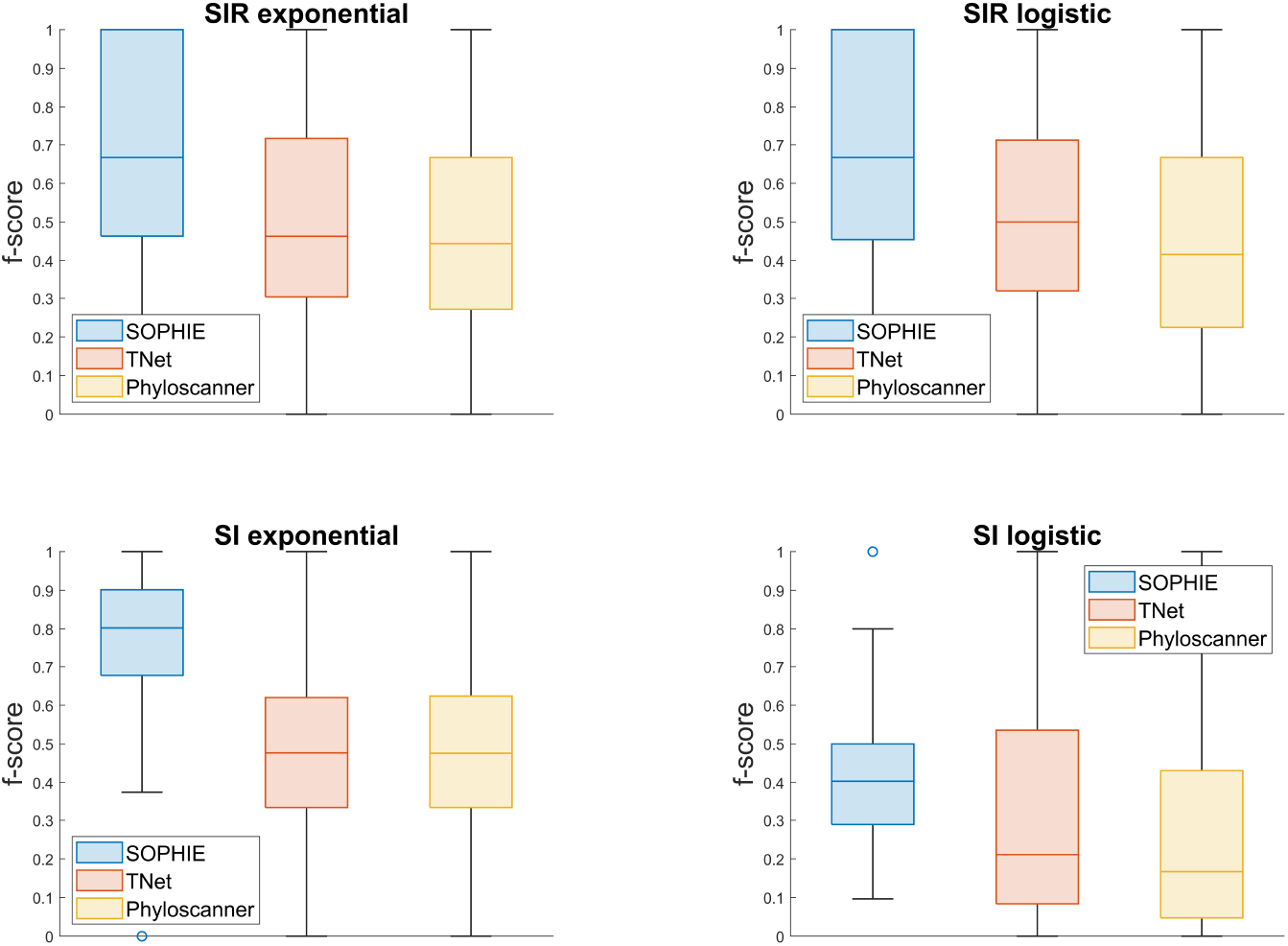
Comparative results of SOPHIE (best exponent), TNet and Phyloscanner on simulated data under different epidemiological and evolutionary scenarios with the tree reconstructed by RAxML

The accuracy of SOPHIE was negatively affected by the phylogenetic inference noise and was generally lower when RAxML tree was used. This effect is less pronounced for TNet and similarly pronounced for Phyloscanner. It is not surprising, since TNet, as a strictly parsimony-based method, depends only on the tree topology, while SOPHIE and Phyloscanner also utilize branch lengths. Still, the accuracy of SOPHIE for RAxML trees remains higher than for other tools.

The results of SOPHIE for different evolutionary and epidemiological scenarios are comparable, with the exception of the Susceptible-Infected transmission model with the logistic intra-host population growth. In that case, the accuracies of all methods were significantly lower.

As described in Methods, all algorithmic subroutines of SOPHIE are polynomial. Thus, the method is not too computationally expensive: the experimental average running time of SOPHIE on the analyzed data was 106.5s (standard deviation 285.4s). It somewhat slower than TNet (with the running times measured in seconds) and Phyloscanner (that stops within 1-2 minutes), but it is to be expected, given that the SOPHIE’s model is richer than for other tools.

#### 2.2.2 Experimental data

We used a “gold standard” experimental dataset that has been previously utilized for benchmarking of transmission network inference algorithms in several studies [67, 30, 21]. It consists of 74 intra-host HCV populations sampled and sequenced during the investigation of 10 outbreaks by the Centers for Disease Control and Prevention. Viral populations contain from several dozen to several hundred sequences of lengths 264bp covering Hypervariable Region 1 (HVR1) of the HCV genome. In each outbreak, a single primary host identified by the investigators using epidemiological evidence infected all other hosts. Thus, transmission networks for that outbreaks are known.

Similarly to simulated data, the algorithms under consideration were applied to phylogenies reconstructed by RAxML. For all outbreaks, the uniform equilibrium label distribution, the rate *μ* = 1 and the power-law exponent *α* = 2 has been used. SOPHIE yielded the average *f* -score of 0.70, while TNet and Phyloscanner showed *f* -scores of 0.58 and 0.37, respectively (Table 1).

### 2.3 Case study: HCV/HIV outbreak in rural Indiana, 2015

We utilized SOPHIE to analyze genomic data from the large HIV/HCV outbreak in Indiana [54, 57, 31, 8, 14]. First 11 HIV infection cases associated with this outbreak have been discovered by the Indiana State Department of Health (ISDH) in a small rural community in Scott County, IN in early 2015. This triggered a further investigation by the ISDH and the CDC [14] that led to detection of several hundred HIV and HCV infections and precipitated a declaration of a public health emergency by the state of Indiana [14]. The investigation linked the outbreak to unsafe injection use of the opioid oxymorphone [54], providing an important example of the rapid spread of viral infections associated with the nationwide epidemic of prescription opioid abuse [80, 71].

Deep sequencing of intra-host viral populations has been carried out only for HCV; therefore, we focused this evaluation on the HCV genomic data. Each HCV dataset consists of viral haplotypes covering the E1/E2 junction of the HCV genome, which contains the hypervariable region 1 (HVR1). We sampled and analyzed transmission networks of the largest HCV transmission cluster identified previously [57]. It includes 116 persons infected with the HCV subtypes 1a and 3a; some persons were infected with both subtypes. The HCV subtypes are phylogenetically distinct. Given that, we first constructed and analyzed maximum likelihood phylogenies for each subtype separately. In addition, these phylogenies were post-processed using TreeTime [62] to infer time labels of their internal nodes. The obtained time-scaled phylogenetic trees were used as inputs of SOPHIE and, after obtaining sampled transmission networks and their probabilities, provided times of inferred transmissions. Finally, transmission networks for both subtypes sampled by SOPHIE were joined into a single network. Further data processing details can be found in Methods section.

The inferred joint consensus transmission network of both subtypes is shown in Fig. 5(a). When reconstructed transmission links have both the person’s metadata and phylogenetic data, they tend to agree with each other. The subcluster of persons infected with subtype 1a is large, established earlier, and is likely to serve as a source for the 3a subcluster. This finding matches the observation that the inferred primary case of the 3a subcluster (the only vertex with the expected in-degree below 1 in the 3a network) is coinfected with both subtypes. In addition, both persons with known acute infection from the analyzed cluster (detected by the HCV seroconversion test) have low expected outdegrees (*<* 10^−4^), confirming that they carried secondary rather than primary infections.

**Figure 5:**
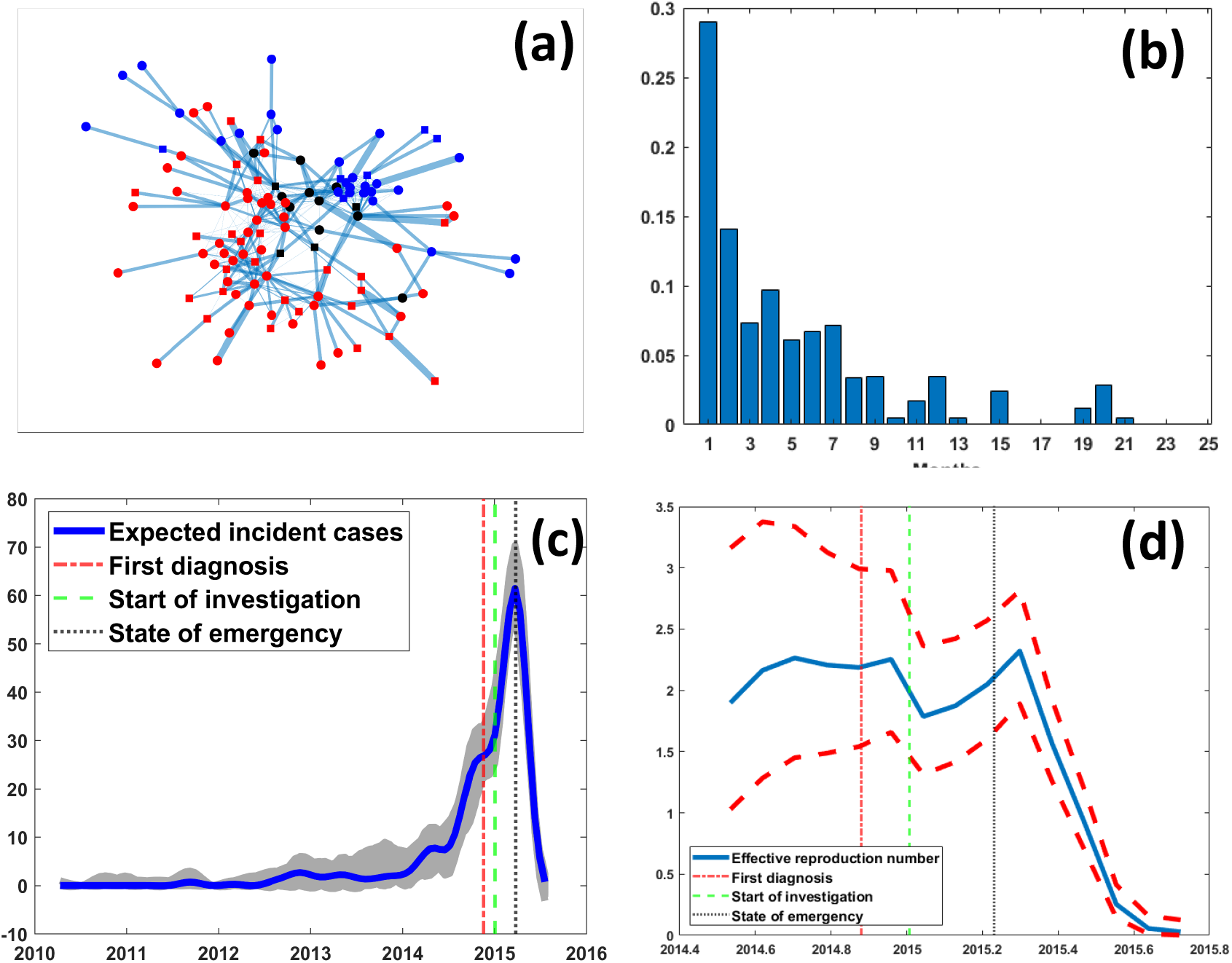
(a) Consensus transmission network of the Indiana HCV outbreak. The thickness of each edge is proportional to its inferred likelihood support. Only edges with the support above 0.0005 are shown. Nodes infected with subtype 1a, 3a and both are shown in red, blue and black, respectively. Squared nodes are coinfected with HIV. (b) Distribution of the generation times by month. (c) The dynamics of incident cases over time. The blue line is the expected number of incident cases at a given time. The grey area shows incident cases for sampled networks. Vertical lines depict major public health events. (d) Effective reproduction numbers *R*_*t*_ for the exponential stage of the outbreak. Vertical lines depict major public health events.

The output from SOPHIE was used to estimate key epidemiological parameters directly from the inferred transmission networks. Such estimates can be more realistic than more traditional assessments based on random mixing models applied to incidence statistics [45]. Furthermore, we used time labels of the viral phylogenies to estimate timing of each link in each sampled transmission network to assess the outbreak dynamics.

The dynamics of incident case numbers (i.e. the numbers of inferred transmissions within a specified time interval, in our case, 1 month) suggest that the outbreak started in the middle of 2012, and transitioned to the exponential stage in 2014 (Fig. 5(c)). The incidence rapidly declined after the declaration of the state public health emergency. The exponential stage largely coincides with the timeline of HIV spread in the same community [8]. In addition, 35 persons from the analyzed cluster were co-infected with HIV, and 25 of them form a connected subgraph of the consensus subnetwork formed by edges with the high support shown in Fig. 5(a). These findings suggest that the HIV outbreak and the larger part of the HCV outbreak were triggered by the same epidemiological mechanism; however; HCV preceded HIV by several years, and the HIV spread might have been facilitated by the pre-established HCV transmission network.

The inferred incidence (Fig. 5(c)) and the inferred distribution of generation times (time intervals between the infection times of the sources and recipients, Fig. 5(b)), were used in EpiEstim [15] to estimate the effective reproduction number *R*_*t*_ (virus transmissibility at a given time) over a 1-month sliding window during the exponential phase of the outbreak. The mean values of *R*_*t*_ varied between 1.81 and 2.33 before the emergency declaration, indicating sustained transmissions. Following the declaration, they rapidly dropped below the epidemic threshold of *R*_*t*_ = 1. We also directly measured the basic reproduction number *R*_0_ as an average degree of transmission sources in sampled networks. An estimation *R*_0_ = 2.71 (95% CI: (2.63, 2.79)) close to the estimates for *R*_*t*_ was obtained. The estimates produced by SOPHIE are more moderate and seemingly more realistic than, for example, the values *R*_0_ = 6.6 (95% CI: (3.2, 9.9)) and *R*_0_ = 5.1 (95% CI: (1.7, 9.2)) produced by the birth-death skyline phylodynamics model [69] with the uniform reproduction number prior implemented in BEAST [26]. Moreover, the SOPHIE-based values agree better with the estimate of *R*_0_ = 3.8 for the parallel HIV outbreak obtained using contact tracing [8].

## 3 Discussion

Analysis of viral transmission networks is essential for epidemiological and evolutionary studies of pathogens, as it allows to assess and monitor the transmission dynamics [2, 5], understand the mechanisms of transmission, infection establishment and emergence of drug resistance and vaccine escape [55, 47], as well as to design efficient public health intervention strategies [11]. Hence, inference of viral transmission networks is one of the most fundamental problem of genomic epidemiology and a major driving force behind new developments in the field.

In this paper, we presented a novel method for transmission network reconstruction based on the integration of a phylogenetic maximum likelihood (ML) model and a random graph model. The idea to implant social networks into the phylogenetic framework was proposed in our prior study [67] and implemented in a new tool, QUENTIN. SOPHIE substantially differs from QUENTIN in several ways: (1) it is fully based on maximum likelihood paradigm, (2) it is phylogenetic rather than network-based, and (3) it uses more general and comprehensive random graph model. In general, SOPHIE re-evaluates phylogeny-based candidate transmission networks according to their match to the expected properties of an unobserved contact network and prioritizes the networks, which fit to both the viral phylogeny and these properties.

We showed that the proposed approach is capable of achieving a substantial accuracy improvement over the state-of-the-art phylogenetic transmission inference methods based on maximum parsimony principle, while retaining their scalability and speed. This improvement is likely associated with the relative sampling efficiency of parsimony and likelihood-based methods. Indeed, for most of the simulated test examples the total numbers of optimal parsimonious solutions (as calculated by TiTUS [65]) were exceedingly large, with the median number of solutions over all tests being 3.9 · 10^15^. Representative uniform sampling from such large set is challenging. In contrast, SOPHIE samples from a more informative distribution. Furthermore, the network-based part of the proposed model allows to optimize the search in the solution space by employing a polynomial-time combinatorial optimization machinery. This distinguishes SOPHIE from other phylodynamics models that are often less computationally tractable and have to rely on the MCMC sampling.

Despite the aforementioned advantages, the proposed methodology has a room for further expansion and improvement. First, its phylogenetic component is currently based on trait substitution models with fixed between-host transmission rates. Incorporation of rate inference via EM or other iterative algorithms can potentially enhance our approach. Such a technique proved to be useful for nucleotide substitution models within traditional phylogenetics and phylodynamics [62]. In our case, however, its application is more challenging due to smaller numbers of ancestral trait changes. Second, ideally the label sampling scheme should simultaneously account for both parts of the joint likelihood. However, use of MCMC or other similar approach for such sampling is non-scalable, while development of the scheme based on combinatorial optimization seems to be challenging. One possible combinatorial approach envisioned by us is the utilization of spectral techniques. Third, as suggested by computational experiments, SOPHIE is sensitive to potential phylogenetic inference inaccuracy, especially, in respect to branch lengths estimation. This can be addressed by allowing for length updates, similarly to transmission rates. Finally, the experiments also revealed the decreased accuracy of SOPHIE (and other methods), when applied to data produced by the SIR transmission model with the intra-host logistic coalescent. This suggests that in this case the method’s accuracy may benefit from replacement of the ML phylogenetic model with the Bayesian coalescent or other appropriate phylodynamic model. Such models are, however, less computationally tractable; therefore their incorporation into our framework will require novel algorithmic solutions.

## 4 Methods

### 4.1 Algorithms

#### 4.1.1 Sampling of ancestral label assignments

Suppose that the Markov chain-based substitution model for labels is fixed, i.e. we are given the equilibrium patient probabilities 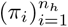 and the rate matrix 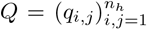, where *q*_*i,j*_ is the transmission rate between hosts *i* and *j* for *i* ≠ *j*, and 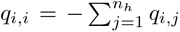. In most cases, transmission rates between specific hosts are unknown. Therefore usually the substitution model will be the fully-symmetric substitution model (similar to Jukes-Cantor model for DNA) with *π*_*i*_ = 1*/n*_*h*_ and *q*_*i,j*_ = *μ/*(*n*_*h*_ − 1), where *μ* is the general transmission rate. In certain cases, however, between-host transmission rates can be assessed from epidemiological contact tracing or comparison of exposure intervals, if such information is available. In that case, more general substitution model can be employed.

Given the substitution model, we sample from the joint distribution of ancestral label assignments using an extension of the Felsenstein pruning [27] - a standard dynamic programming algorithm for phylogeny likelihood calculation. It is a dynamic programming algorithm that performs a post-order traversal of the phylogeny *T* and computes, at each node *v* ∈ *V* (*T*) and for each host *i* ∈ [*n*_*h*_], the conditional likelihood *L*(*v, i*) of observing the labels of leafs that are descendants of *v*, given that *λ*_*v*_ = *i*. The computations are based on the following recurrent relation [27]:

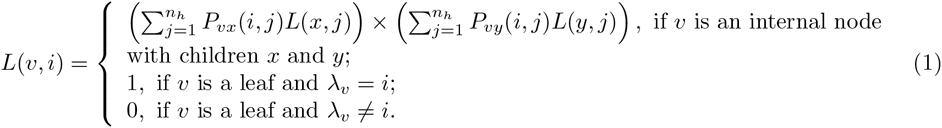

Here *P*_*vx*_ = *exp*(*t*_*vx*_*Q*) is an *n*_*h*_ × *n*_*h*_ transition matrix for an edge *vx* ∈ *E*(*T*), where *t*_*vx*_ is the length of *vx*.

##### Algorithm 1 Ancestral label sampling

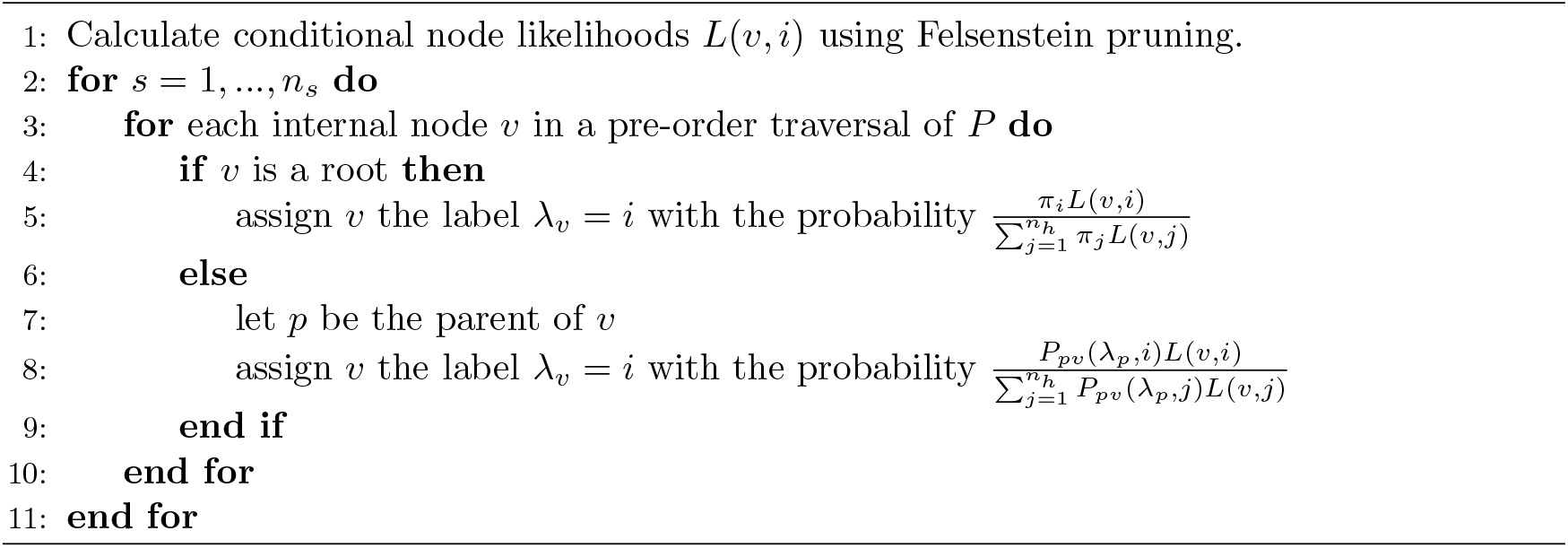

After all conditional likelihoods *L* are calculated, we perform a pre-order traversal of *T* and sample a label for each node from the corresponding posterior distribution given its parent’s sampled state. The sampling is repeated *n*_*s*_ times. The sampling procedure is formally described by Algorithm 1. For each sampled label assignment ***λ*** = (*λ*_*v*_)_*v*∈*V* (*T*)_, its *phylogenetic likelihood L*(*T*, ***λ***) is calculated as 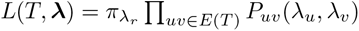, where *r* is the root of *T*.

For large phylogenies, the number of ancestral label assignments with comparable likelihoods can be large. Thus, in order to facilitate sampling of the assignments that potentially produce transmission networks with high network likelihoods, we employ several heuristic adjustments of the general sampling scheme. First, we reduce the tree before sampling by iteratively removing sibling leafs with the same label and assigning that label to their parent. This procedure replaces all monophyletic clades with their most recent common ancestor. This modification decreases the dimensionality of the ancestral label space, thus allowing to obtain a representative sample with fewer iterations. In addition, it speeds up likelihood calculations and decreases the number of likelihood re-scalings [77] required to resolve the numerical precision issues. Next, it is known that intra-host viral population diversity can serve as a marker of the population age [4], and therefore hosts with more diverse populations are more likely to be sources of transmissions [67, 76, 61]. We account for that by multiplying the likelihoods *L*(*v, i*) calculated for the reduced tree by the number of descendants of *v* with the label *i*.

The total running time of the sampling step is 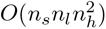

#### 4.1.2 Estimation of the network likelihood

##### Likelihood definition

We assume that the transmission network *G* = *G*(*T*, ***λ***) is a subgraph (not necessarily induced) of a random contact network 𝒢_*c*_ on *n*_*c*_ ≥ *n*_*h*_ vertices. We model G_*c*_ as a random graph with the given degree distribution ***p*** = (*p*_1_, *p*_2_, …), where *p*_*k*_ is the probability that a randomly selected vertex has a degree *k*.

Every vertex *i* ∈ *V* (*G*) has a degree *d*_*i*_ in *G* and a degree *D*_*i*_ ≥ *d*_*i*_ in 𝒢_*c*_. Let us call a mapping 𝒟 : *V* (*G*) → [*n*_*c*_ − 1], that assigns a degree 𝒟(*i*) = *D*_*i*_ to a vertex *i*, an *embedding of G into* 𝒢_*c*_. Then we approximate the network likelihood *L*(*G*|𝒢_*c*_) via the probability of the best embedding:

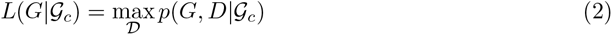

To define the conditional probability *p*(*G*, 𝒟| 𝒢_*c*_), we can factorize it as

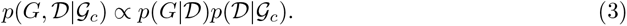

The first factor *p*(*G*|𝒟) is the probability of the subgraph *G* given the degrees of its vertices in the contact network 𝒢_*c*_. It can be calculated by assuming that *n*_*c*_ is large enough and 𝒢_*c*_ follows assigned to pairs of vertices (*i, j*) with probabilities 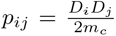, where 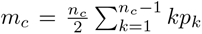 is the *Generalized Random Graph (GRG) model* [12, 13] – a general and widely used model of a random graph with given expected degrees. According to this model, edges are independently the expected number of edges of 𝒢_*C*_. Using this definition, we get

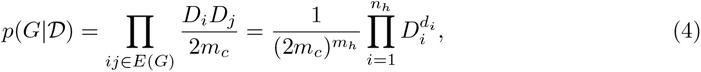

where *m*_*h*_ is the number of edges of *G*.

To define the second factor *p*(𝒟|𝒢_*c*_), consider the vector of expected degree counts 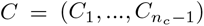 of 𝒢_*c*_, i.e. *C*_*j*_ = ⌈*p*_*j*_*n*_*c*_⌉ is a rounded expected number of vertices of degree *j*. Then *p*(𝒟| 𝒢_*c*_) is the probability that the degrees 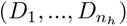 are sampled without replacement from the population *C*. Thus, *p*(𝒟|𝒢_*c*_) is described by the probability mass function of the multivariate hypergeometric distribution:

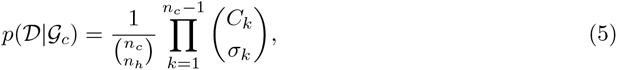

where *σ*_*k*_ = | 𝒟^−1^(*k*)| = |{*i* : *D*_*i*_ = *k*}|.

##### Likelihood calculation

To calculate the network likelihood, we need to solve the optimization problem (2). After logarithmic transformation, it is equivalent to the following problem:

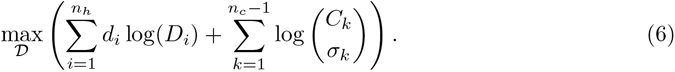

In turn, this problem can be reduced to a *generalized uncapacitated facility location problem with convex costs* [25], where the vertices of *G* serve as clients and their possible expected degrees in 𝒢_*c*_ – as facilities. More specifically, we consider the set of clients *K* = [*n*_*h*_] and the set of facilities *F* = [*n*_*c*_ − 1]; if the client *i* is served by the facility *k* (i.e. *D*_*i*_ = *k*), where *k* ≥ *d*_*i*_, then the profit *b*_*ik*_ = *d*_*i*_ log(*k*) is generated. Furthermore, the assignment of *σ*_*k*_ clients to a facility *k* produces a profit 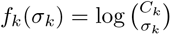. The objective is to assign all clients to facilities in such a way that the total profit is maximized.

The crucial property of the obtained problem is the fact that the functions *f*_*k*_(*σ*) are concave (or, if we are using more standard minimization formulation, − *f*_*k*_(*σ*) are convex). Thus, we can use the scheme proposed in [33] to reduce our problem to the maximum-weight matching problem for bipartite graphs, which is solvable in polynomial time [66]. Namely, we construct a bipartite graph *H* with the parts (*X, Y*), where the part *X* coincides with the set of clients *K*, and the part *Y* contains *C*_*k*_ vertices 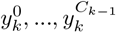 for each facility *k*. The vertices *i* ∈ *X* and 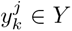 are adjacent whenever *d*_*i*_ ≤ *k*, and the weight of this edge is set to 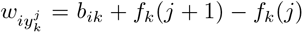. Then maximum-weight matching of *H* gives us the solution of (6). This fact follows from the concavity of the function *f*_*k*_, which implies that any maximum-weight matching that covers the vertex 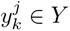 should also cover all vertices 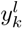 for *l* ≤ *j*.

It is easy to see that the number of edges in the bipartite graph *H* is *n*_*h*_(*n*_*c*_ + 1) 2*m*_*h*_. Therefore, the described reduction approach combined with the generalized Hungarian algorithm for the matching problem [58] calculates the network likelihood in time 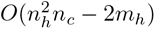.

Finally, it should be noted that the model (3) contains the size of the contact network *n*_*c*_ as a parameter. In our calculations, we used the value that is large enough to guarantee the existence of a feasible solution of (6), i.e. *n*_*c*_ = max_*i*_⌈*c*_*i*_*/p*_*i*_⌉, where *c*_*i*_ = |{*j* : *d*_*j*_ = *i*}| are degree counts of *G*. In particular, if the expected degree distribution of 𝒢_*c*_ follows the power law with the exponent *α*, then *n*_*c*_ can be estimated as 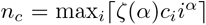, where *ζ*(*α*) is the Riemann zeta function.

#### 4.1.3 Distribution and consensus of sampled networks

The output of the algorithms described above is the set of *N* sampled solutions, where each solution consists of the label assignment ***λ***^*i*^, the corresponding transmission network *G*(*T*, ***λ***^*i*^) and the joint likelihood *L*(*T*, ***λ***^*i*^)*L*(*G*(*T*, ***λ***^*i*^) _*c*_). The distributions of transmission networks and labels, as well as derivative epidemiological parameters, can be further analyzed directly – an example of such analysis for a particular case study is presented in Subsection 2.3. In particular, sampled networks can be summarized into the weighted *consensus network* with the adjacency matrix 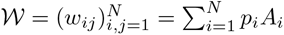, where *A*_*i*_ is the adjacency matrix of the network *G*(*T*, ***λ***^*i*^), and 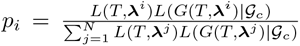 is the probability density value estimate for that network. In this case, *w*_*ij*_ is an inferred likelihood support for an edge *ij*, and 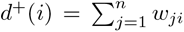 and 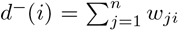 are expected in- and out-degrees of a vertex *i*, respectively. When a specific output network is needed (e.g. for benchmarking, see Subsections 2.2.1-2.2.2), then we calculate it as the maximum-weight arborescence of this weighted network.

### 4.2 Quantification and statistical analysis

#### 4.2.1 Simulation and algorithm comparison details

Synthetic data used in this study was generated by FAVITES [50]. Viral genomes of length 2640bp (that roughly corresponding to lengths of HIV gap and pol polyproteins) were assumed to evolve under the GTR+Γ substitution model. The GTR rate matrix and gamma parameter were borrowed from [68], where they were estimated based on real HCV data. Inside each host, viral phylogenies evolved under a coalescent model with exponential or logistic effective population growth. We assumed that the virus spread over a contact network of 100 susceptible individuals, that was produced using the Barabasi-Albert model [3]. Two epidemiological scenarios were used: Susceptible-Infected (SI) transmission model and simultaneous sampling of all infected individuals and Susceptible-Infected-Recovered (SIR) transmission model, with each individual sampling time being chosen from a truncated normal distribution of the individual’s infection time window. The full lists of FAVITES parameters are available in configuration files provided with simulated datasets in SOPHIE repository.

For each of four combinations of evolutionary and epidemiological models, 100 simulated datasets have been generated, with 10 genomes sampled per infected host. Simulations that produced no transmission links were discarded. For each dataset, we considered a true phylogeny provided by FAVITES and a phylogeny reconstructed by RAxML [70]. The latter was run with the GTR+Γ substitution model, and with optimization of substitution rates and site - specific evolutionary rates.

TNet was run with the default settings. For Phyloscanner, we set the within-host penalty parameter to 0 (otherwise, it produced no transmission links). For SOPHIE, at the label sampling stage we used the uniform equilibrium probability distribution and fixed transmission rates *μ* = 0.0001 and *μ* = 0.005 for all Favites and RAxML trees, respectively. For each test instance, 100, 000 internal label assignments were sampled. The *f* -score has been used as an evaluation metric. To compare the distributions of *f* -scores for different algorithms, we utilized a nonparametric Kruskal–Wallis test.

### 4.2.2 Analysis of HCV outbreak in rural Indiana

Analyzed HCV datasets consist of viral haplotypes sampled from infected individuals and sequenced using GS FLX Titanium Sequencing Kit (454 Life Sciences, Roche, Branford, CT). The haplotypes cover the E1/E2 junction of the HCV genome (264 bp), which contains the hypervariable region 1 (HVR1). For our analysis, we used haplotypes that were sampled at least 5 times in each infected person. In total, 4167 viral haplotypes (or ≈ 36 haplotypes per person) have been considered. Prior to phylogenetic analysis, the sequences have been aligned using MAFFT [40]. Next, maximum likelihood phylogenies were constructed for each subtype; in addition, these phylogenies were time-labelled using TreeTime [62] run with default parameters. The obtained time-scaled phylogenetic trees were processed by SOPHIE, for which we used the uniform equilibrium label distribution, the rate *μ* = 1 and the power-law exponent *α* = 2. For each phylogeny, 2, 000, 000 label assignments were sampled.

## 5 Acknowledgements

PS was supported by the NIH grant 1R01EB025022 and by the NSF grant 2047828. VT was supported by the GSU MBD Fellowship. The authors thank M. Bansal, P. Sashittal, M. El-Kebir and M. Hall for their help in running TNet, TiTUS and Phyloscanner.

## 6 Disclaimer

The findings and conclusions in this report do not necessarily reflect the official position of the Centers for Disease Control and Prevention, or the authors’ affiliated institutions.

## 7 Ethical approval

The Centers for Disease Control and Prevention approved this secondary research with existing outbreak data.

## 8 Code availability

SOPHIE code is freely available at https://github.com/compbel/SOPHIE/

